# An elementary model of homeostasis and immunity that generates symbiosis

**DOI:** 10.64898/2026.01.15.695950

**Authors:** Gérard Eberl

## Abstract

The immune system was historically defined as a system that provides protection from pathogens. Numerous models have been developed to understand how immunity faces a complex world of microbes that includes pathogens and symbionts, as well as cells of our own self that may develop tumors. Based on the classical assumption that survival depends on internal homeostasis, we have developed a formal model of homeostasis for a host interacting with microbes and self. We propose that such a model must include two fundamental functions: a function that counters change (including tissue repair), and a function that counters the agent of change (such as “immunity” to microbes or self). We show that this elementary model is sufficient to generate symbiosis, and that symbiosis is an emergent property of the host-microbe relationship that does not require the microbe or the host to express “traits of symbiosis”. We suggest that the conditions leading to symbiosis contribute to eukaryotic evolution and ontogeny. This model may be further applied to symbiotic interactions between organisms and non-microbial or non-cellular agents of change.

## PATHOGENS AND SYMBIONTS

How does the immune system discriminate a pathogen from a symbiont? This question, which we would like to challenge here, has become preeminent ever since immunologists realized the extent of the crosstalk between the symbiotic microbiota and the immune system ^1-6^. For example, the large intestinal microbiota, far from being walled off and ignored, induces robust immune responses that are necessary to maintain a vital equilibrium with the host. These responses include the production of mucus and anti-bacterial peptides by intestinal epithelial cells, IgA antibodies that restrict the access of microbes to these epithelial cells, the production of cytokines and chemokines by hematopoietic and non-hematopoietic cells, and the recruitment and activation of a wide array of lymphoid and myeloid cells. This regulated state of inflammation, which is part of “business as usual”, has been termed “physiological inflammation” or “homeostatic inflammation” ^4,7,8^.

Upon infection by pathogens, this regulated state of physiological inflammation transits to so-called “pathological inflammation” that is toxic for the microbes (and the host). Furthermore, deregulation of inflammation (for example in patients and mice that are deficient in potent regulators of immunity, such as interleukin-10) leads physiological inflammation to become pathological and reveal the “concealed” pathogenic potential of certain members of the microbiota, such as *Helicobacter hepaticus*, which are thereby termed “pathobionts” ^9-11^.

Many mechanisms have been proposed that may allow the host to distinguish a pathogen from a symbiont, such as the expression of virulence factors by microbes that are recognized directly by the immune system, the activation of bait proteins or pattern recognition receptors (PRRs) that signal the presence of pathogens, and the penetration of microbes into “forbidden” spaces, such as the cytoplasm ^12,13^. Conversely, it has been proposed that the immune system recognizes positive microbial traits, sometimes termed symbiosis factors ^14^, or that symbionts activate select types of immune cells that induce immunological tolerance ^15-17^.

These mechanisms are based on the proposition that pathogens and symbionts differ fundamentally in form or function. In contrast to this hypothesis, it has been proposed that a microbe is not intrinsically pathogenic or symbiotic, but rather that the host-microbe relationship can be pathogenic or symbiotic ^18-20^. In support of this view, microbes may benefit the host in one context, such as providing increased resistance to superinfection, but threaten the same host in another ^21-24^. This “relativistic” view of microbes is interesting, because it suggests the existence of a simpler set of rules to describe the individual host-microbe relationship in a continuum between pathogenicity and symbiosis ^25^.

Here, we have developed a formal model of pathogenicity and symbiosis that is based primarily on the state of the host. Our assumption is that the host’s fundamental physiological aim is to maintain internal homeostasis ^26-28^, and thus, to counter any change induced by any type of stressors (such as microbes, parasites, tumor cells, wounds, physicochemical conditions, …). To fulfill this aim, we argue that the host requires two functions: one to directly counter change (the homeostatic function), and another to counter the agent of change (the resistance function). Surprisingly, these two functions lead, together, to new states of homeostasis (or “homeostates”) characterized by the integration of the agent, or symbiosis, as well as to host evolution and complex ontogeny.

## HOMEOSTASIS AND ITS FORMALIZATION

The term *homeostasis* has been coined in 1926 by Walter Cannon ^27^, but the concept that the physical, chemical and biological variables of an organism must be kept within a range of values to maintain life has been articulated by Claude Bernard in 1849 ^26^. Bernard’s modern physiology may reflect, in more realistic terms, Hippocrates’ (5^th^ century BC) and Galien’s (2^nd^ century AD) antic notion of a necessary equilibrium between four fundamental body humors (blood, yellow bile, black bile and phlegm), failure of which would drive pathology. The common principle is that life forms have to maintain an internal equilibrium in the face of external (and internal) perturbations in order to survive and perpetuate.

For example, blood pressure, glycemia, pH or oxygen levels have to be maintained within a vital range of values. This principle is true at all levels, from organs to tissues, cells and molecules, which have to be constantly monitored and corrected, and our genome, which should suffer minimal damage and be replicated with minimal errors to avoid deleterious mutations. As discussed in detail previously ^28^, mechanisms of sensing and correction (by repair and regeneration) are in place at every level to maintain homeostasis. Nonetheless, aggressive agents may inflict too much damage for such processes to be able to maintain homeostasis. Therefore, additional “resistance” processes may be necessary to directly target agents and prevent them from inflicting more damage and irreversibly affect homeostasis.

Numerous mathematical models have been developed to characterize, for example, glucose homeostasis ^29,30^, based on the knowledge of the sensors and correcting effectors involved in the process. Here, we propose a general model of homeostasis that includes both a “corrective” function and a “resistance” function, and apply the model to host-microbe interactions. We will first describe, as briefly but comprehensively as possible, the mathematical foundations of the model, before discussing its main predictions for symbiosis.

In a host that perfectly enforces homeostasis, any deviation D from homeostasis is immediately corrected (D = 0). However, more realistically, an agent of change A (such as a microbe) will induce at least transient deviation in the host, which can be written as:

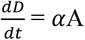

(where a change in D is induced by the agent A at a rate 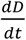 that depends on the quantity of agent and its impact *α* on the host). In order to maintain or restore homeostasis, a homeostatic function (H) is required to counter that change and bring D back to zero. This can be written as:

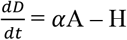

(no Greek letter to H here, because, as we will see below, that letter is included in the description of H). However, as we mentioned above, this may not be enough to maintain homeostasis if the agent is numerous, replicating fast and its impact *α* on the host is high, thus creating deviation faster than the homeostatic function can cope with. Therefore, a resistance function R must be evoked that directly targets the agent, a function that intuitively includes our notion of immunity. Since it directly targets the agent, the resistance function decreases D indirectly, but also increases D by affecting the host tissues and resources (as immune responses do, in general, by a factor *β* that is at best zero: *β* ≥ 0). So, we propose the following equation to be the general expression of homeostasis:

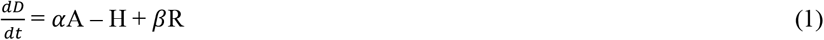

In our view, this is the most general formalization of homeostasis in living systems, which may be applied to any system of cells, multicellular organisms, groups of individuals and ecosystems once the terms A, H and R are defined in these contexts. In the next section, we will apply this general equation to the host-microbe relationship.

## THE HOST-MICROBE RELATIONSHIP

Let us first formally define the microbial agent A. In its simplest form, the expansion of a microbe (be it a virus, a bacteria or a protist) follows an exponential curve as a consequence of replication, which is eventually curbed by an upper limit (an asymptote) as resources and space are limited (this defines the niche of the microbe in the host). The expansion of the microbe in the host is thus described by the “logistic” function 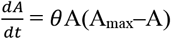, where A_max_ is the maximum possible number of microbes in the niche, and *θ* is the intrinsic growth rate of the microbe. However, the microbial agent faces the host’s resistance function R, and so its expansion in the host is more accurately described as:

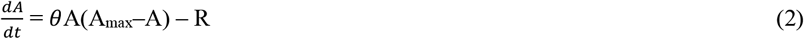

Let us now define the homeostatic (H) and resistance (R) functions. As the homeostatic function H directly reacts to deviation from homeostasis, and the resistance function R directly reacts to the agent of change, the simplest proposition is that these functions are proportional to the level of deviation and agent, respectively (other formulations are discussed in Figure S2):

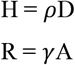

However, these mathematical descriptions of the host’s functions do not yet match the real world, for two reasons. The first reason is that reactions in the host take time and energy, while these mathematical descriptions do not include boundaries in time and energy. The homeostatic function needs time to repair a tissue, and the resistance function needs time to unfold against an agent (these functions typically follow logistic functions as they involve self-amplifying molecular and cellular reactions). Furthermore, the host’s resources are limited, and the host’s functions can exhaust these resources (e.g. the fatigue induced by the immune response against an infection) and compete against each other (e.g. repair responses competing with immune responses) ^24^. We thus introduce into our formalization the mathematical function Ψ that reflects these limitations (Figure S1).

The second reason that these mathematical descriptions do not yet match the real world is that the resistance function (R) reacts to an agent of change (A) that (by definition) induces a deviation. Our current formulation, R = *γ*A, implies however that the host can react to an agent (A) that has no impact on the host. This issue is corrected by making R also reactive to the change induced by the agent, or more generally, to the global deviation induced by the microbe-host relationship (R ≈ δD). Of note, even though it may appear intuitive, from the immunological point of view, that the host reacts to any microbial agent recognized as “non-self” in order to prevent the microbe’s infection of the host, observations reported in the 1980’s and 1990’s show that antigens delivered without some form of tissue damage or adjuvants, do not elicit an immune response, but rather antigen-specific lymphocyte anergy ^31-35^. Pattern recognition receptors (PRR) ^36^, which will be discussed in different contexts below, measure deviation via the recognition of microbe- (MAMPs) and damage- (DAMPs) associated molecules ^12,37,38^, as well via interaction with neighboring cells ^39,40^, and provide such adjuvant effects. Thus, the homeostatic and resistance functions are more accurately formalized as follows (and multiplied by Ψ if studying early reactions in the host-microbe relationship, or in situations of limited resources, Figure S1):

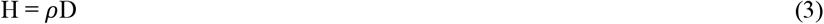

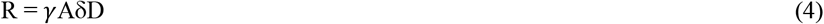

Equations 1-4 describe the dynamic of the host-microbe relationship (Figure S2). We provide the R code for the resolution and simulation of this dynamic (Figure S3), so the readers can explore the model. Its outputs are the level of agent A in the host, the host’s deviation D from homeostasis, and the activation levels of the host’s homeostatic H and resistance R functions (Figure 1). The model has a mathematical limit at 1, meaning that the values for the host variables (D, H and R) cannot equal or exceed 1. This limit can be understood as the maximal available resources in the host, which cannot be reached without death of the host. We close here the mathematical description of the model, for which additional elements and alternative formulations are discussed in figure S2.

**FIGURE 1.**
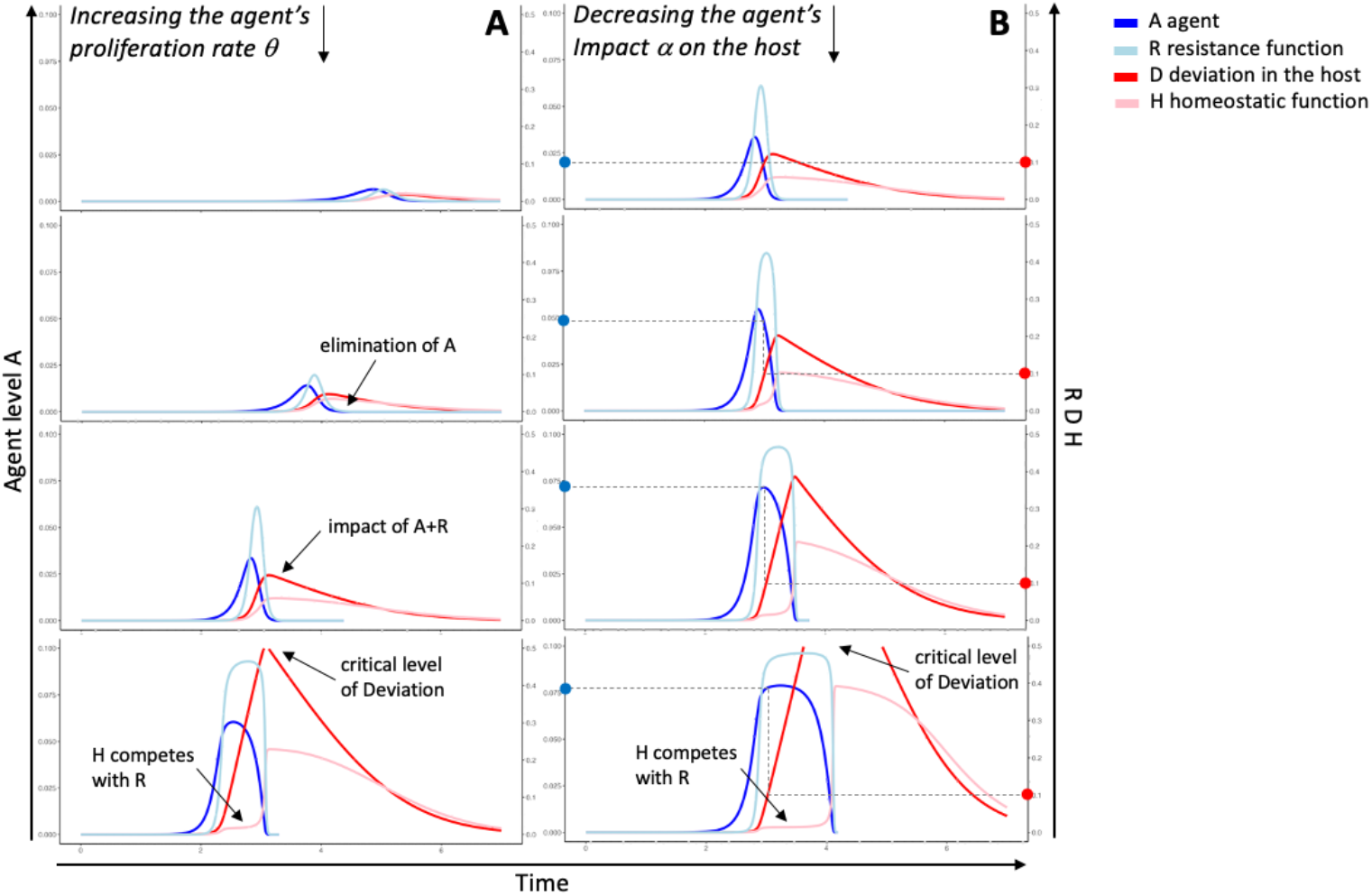
The agent’s impact on the host-microbe relationship. The evolution in time of the host-microbe relationship with the microbe’s (**A**) increasing intrinsic proliferation rate and (**B**) decreasing impact (as deviation) on the host. Shown are the levels of agent (A), the host’s deviation from homeostasis (D), homeostatic function (H) and the resistance function (R). In b), dots on the axes show that increasing amounts of agent (blue) with decreasing impact induce the same level of deviation to the host (red). In all depicted cases, the agent is eliminated from the host, or the host is close to death as deviation reaches critical levels. Note that deviation is induced both by the agent and the host’s resistance function (impact of A+R), and that the homeostatic function competes with the resistance function when the host’s resources become limited (H competes with R). These graphs have been generated by the resolution and simulation of the systems of equation describing the host-microbe relationship (Figure S2) by R (Figure S3). Parameter values: α=1.5; β=1.4; γ=25; δ=35; ε=0.6; τ=5.0; ρ=2.0; δ=3.5; A_max_=1.8. In a) δ = 1.86; 2.5; 3.5 and 4.27. In b) α = 1.5; 0.5; 0.25 and 0.21. Resources parameters: ω=1, ν=10, ah=0, bh=10, ch=0, dh=0.

Figure 1A shows the response of the host to microbes with different proliferation rates (θ), from slow (as would be expected from an innocuous microbe) to fast (as would be expected from a virulent microbe). In these conditions, “slow” microbes are eliminated from the host, while “fast” microbes rapidly overwhelm the host’s resources for its homeostatic and resistance functions, and thus lead to lethal deviation from homeostasis. Figure 1B shows the less trivial reaction of the host to microbes with different impacts (α), from a microbe with high impact on the host’s deviation (damaging microbe) to a microbe with low impact on the host’s deviation (stealthy microbe). A low-impact microbe will expand more than a high-impact microbe (blue points on left y-axis) to induce the same effect on the host (red points on right y-axis). As a consequence, the host’s responses will be induced late in the microbe’s expansion, at levels that are eventually higher, competing for the host’s resources, and potentially unable to control critical levels of deviation.

The situation described in Figure 1B may reflect divergent individual outcomes to an infectious microbe, such as to SARS-CoV-2. The “devious” presence of significant levels of viral RNA in a cell compartment that does not normally carry such RNA ^13^ may be detected early by PRRs and induce protective type I interferon production. In contrast, failure to detect such early deviations, because of less RNA production by the virus in that compartment, or less efficient sensing of such local deviations by the host, may eventually lead to a damaging (and potentially lethal) cytokine storm induced by higher levels of virus and tissue damage ^41,42^.

It is important to note that our model is fundamentally devoid of “valence”. An agent of change, by definition, induces a deviation to the host. Whether this deviation is beneficial or detrimental to the host is determined at a different level, as are the eventual effects of the host’s resistance and homeostatic functions. We will discuss these points in the next chapters.

## MICROBIAL SYMBIOSIS

Figure 1 shows host-microbe relationships that lead to the elimination of the microbe from the host, or to the host’s eventual death. It also shows that the deviation to the host that is induced by the microbe (in the form of tissue damage, for example), can remain significant “long” after the microbe has been eliminated, as the host’s homeostatic function requires time to reduce the deviation back to zero (repairing damaged tissues, for example). It seems obvious that the host would seek to increase the reactivity of the homeostatic response to minimize the time to revert to normality. Figure 2A shows the effect, on the host-microbe relationship, of an increase in the reactivity of the homeostatic function. At first, as expected, an increase in the reactivity of the homeostatic function leads to a faster resolution of the deviation. However, surprisingly, a further increase in the reactivity of the homeostatic function leads to a resurgence of the microbial agent, which is subsequently controlled by a transient increase in the resistance function. This cycle of microbial control and resurgence repeats itself, dampening with time and oscillating around equilibrium values for the levels of microbes in the host, host deviation and homeostatic and resistance functions required to maintain this equilibrium. However, if the reactivity of the homeostatic function is increased further (when the host literally “forgets” about the effects of the microbe, erasing memory to it, encoded as tissue damage, “sequelae” or immunological memory), the transiently dampened deviation leads to lower levels of resistance, increased microbe expansion, and eventually lethal deviation levels.

**FIGURE 2.**
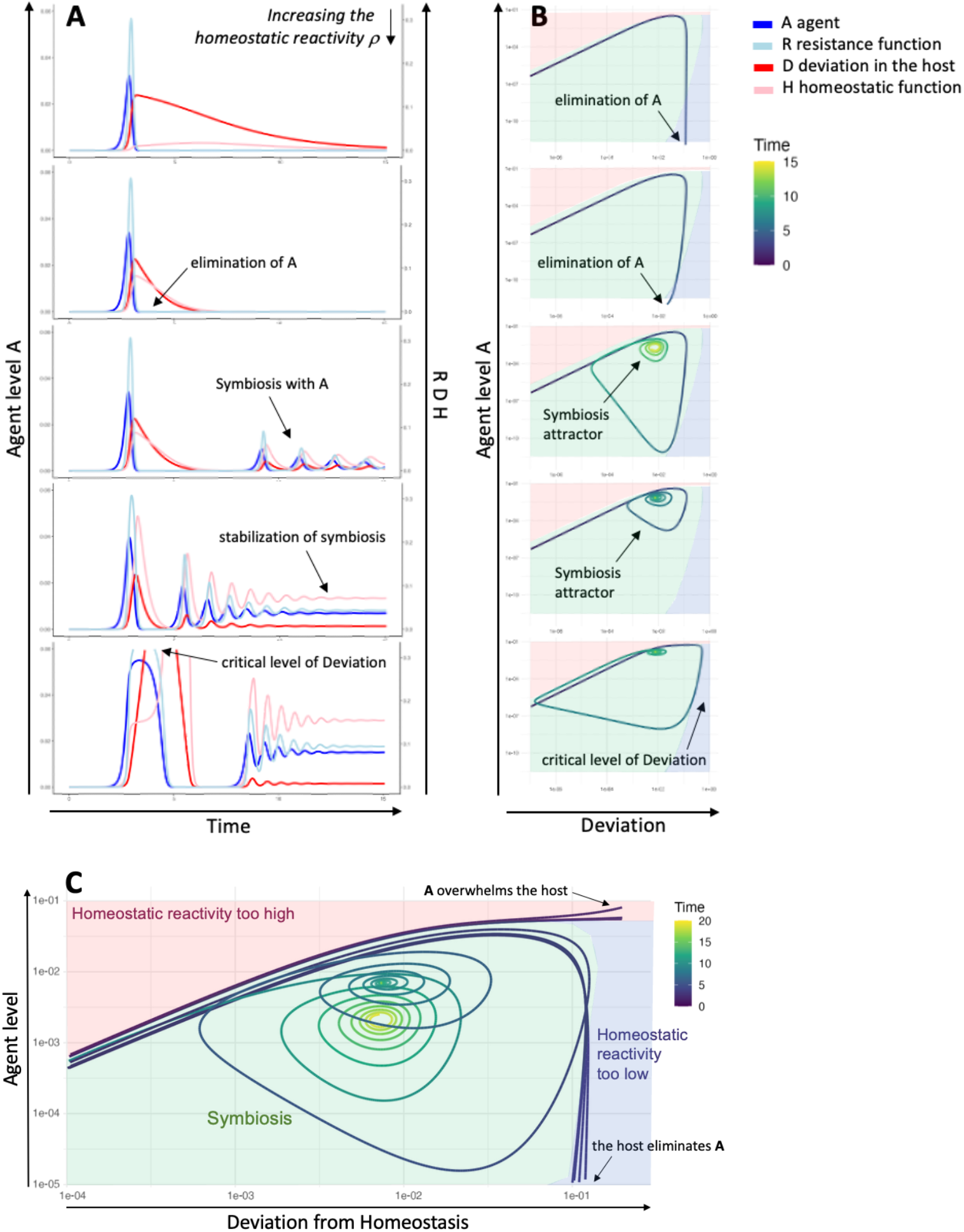
The generation of symbiosis. The evolution in time of the host-microbe relationship with the host’s increasing levels of homeostatic function. At some level of the homeostatic function (H), the agent is no more eliminated, and symbiosis is established between the host and the microbe. However, at higher levels of H, the host is close to death as deviation reaches critical levels, as if the host was “forgetting” about the agent’s effects. **A**) Shown are the levels of agent (A), the host’s deviation from homeostasis (D), homeostatic function (H) and the resistance function (R). Parameter values are as in Figure 1, except for ρ = 0.31, 2.8, 3.02, 10 and 21.66. **B**) Phase diagrams of the evolution in time of the host-microbe relationship, as a function of A and D, with the homeostatic reactivity corresponding to the graphs in a). Shown in green is the attraction basin of symbiosis to the eventual point of equilibrium (the symbiosis attractor), in red the “escape” basin of A overwhelming the host lethally, and in blue the escape basin of the host eliminating A. **C**) Phase diagrams for ρ = 1, 2, 2.8, 2.8056, 3, 10, 21.7232, 21.7235, 22, 25), with all other parameters as in a).

Importantly, Figure 2A describes the *emergence* of a symbiotic host-microbe relationship. The host and the microbe establish an equilibrium that, paradoxically, requires the “anti-change” homeostatic function of the host, as well as its “anti-agent of change” resistance function, to be maintained (Figure 3). In that line of ideas, it has been proposed that border (mucosal and skin) tissues harbor a high level of homeostatic functions, such as repair and regeneration, in order to “tolerate” high levels of microbes ^43^, while the high level of specificity provided by adaptive immunity in vertebrates allows for efficient microbe control and, as a consequence, for the establishment of complex symbioses ^44^.

**FIGURE 3.**
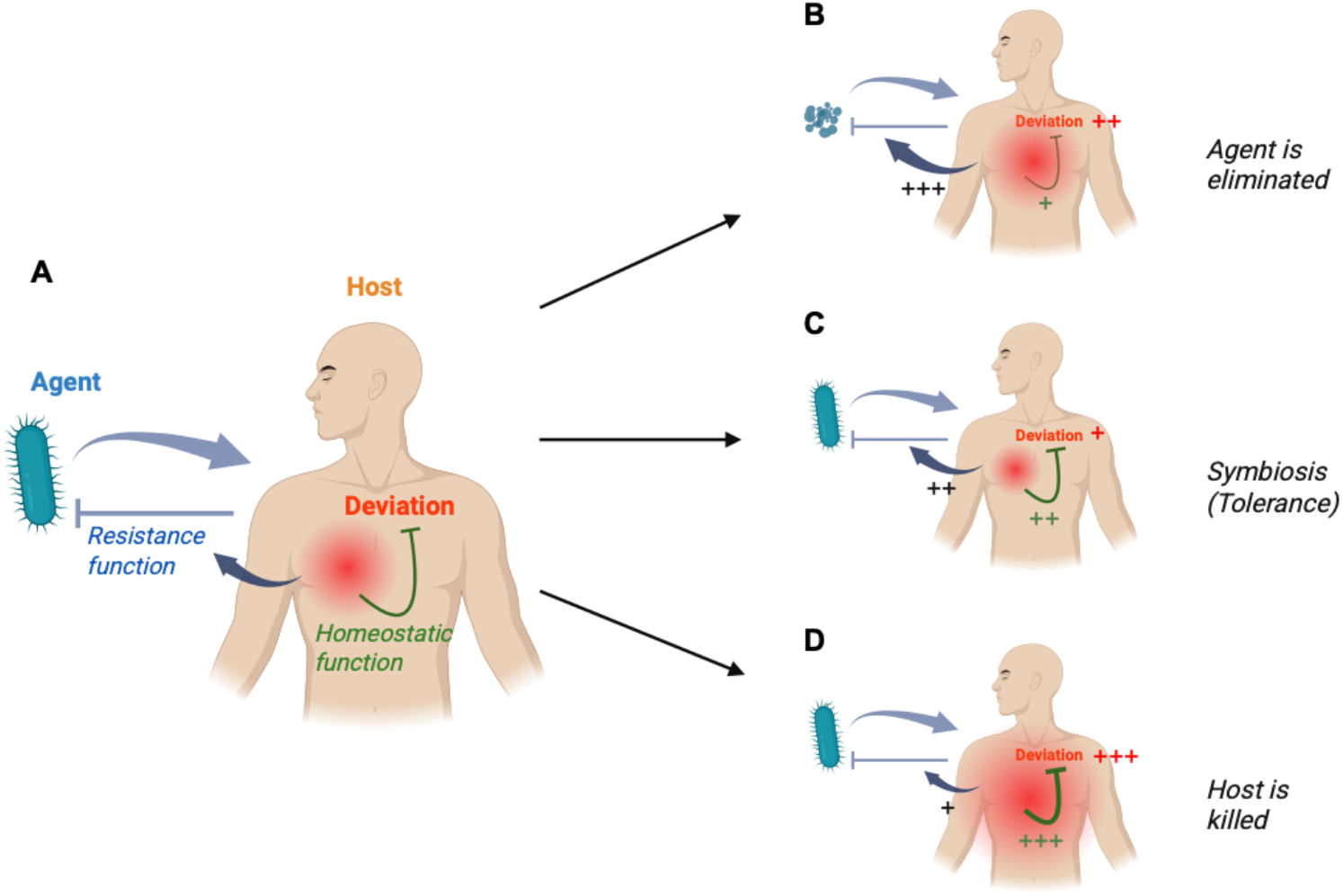
Symbiosis as a consequence of homeostasis and resistance. The model presented here is based on the assumption that homeostasis is key for life. **A**) When an agent of change, such as a microbe, interacts with a host, the host reacts with a homeostatic function to reduce any deviation from homeostasis that is induced by the agent. However, this function may be overwhelmed if the agent is highly virulent and damaging, unless a resistance function directly targets the agent of change. **B**) If the microbe is inducing significant deviation (such as damage) to the host, a significant resistance function is induced that eliminates the microbe from the host. **C**) If deviation is reduced enough by the homeostatic function, the resistance function decreases, and a state of symbiosis is established. **D**) However, a further increase in the homeostatic function decreases the resistance function to levels unable to control for the microbe, eventually leading to lethal deviation in the host (See Figure 2).

The generation and limits of symbiosis can be visualized as phase diagrams – the evolution in time (trajectories) – of the host-microbe relationship (Figure 2C): If the homeostatic reactivity is too high, the host-microbe relationship evolves to a point where the microbe eventually overwhelms the host, whereas if the homeostatic reactivity is too low, the relationship evolves to a point where the host eventually eliminates the microbe; within these limits, the host-microbe relationship evolves to symbiosis, visualized as an attraction basin towards an eventual point of equilibrium. Any host-microbe trajectory that exits the attraction basin leads to a loss in symbiosis (Figures 2B, C). Importantly, the limits of symbiosis follow the chaos theory ^45,46^: near these limits, a small change in the conditions of the host-microbe relationship has a dramatic effect on the outcome of this relationship, i.e. microbe elimination or host death. Thus, even though symbiosis and death are clearly different outcomes, the difference between a symbiotic and a lethal relationship can be infinitesimally small for particular host-microbe relationships ^47,48^. Furthermore, during symbiosis, the microbe is stably integrated into the host, as long as the host’s resources and functions, as well as the microbe’s parameters, remain unchanged (see the conditions of symbiosis, Figure S4). If for example the host’s resources diminish as a consequence of injury, disease, aging or stress, this equilibrium is perturbed and may evolve to the loss of the microbe, or pathology and death of the host (Figure 4). Similarly, a change in the microbe’s parameters, such as its proliferation rate or niche in the host, may turn a symbiotic relationship into a pathogenic and lethal relationship (Figure S5).

**FIGURE 4.**
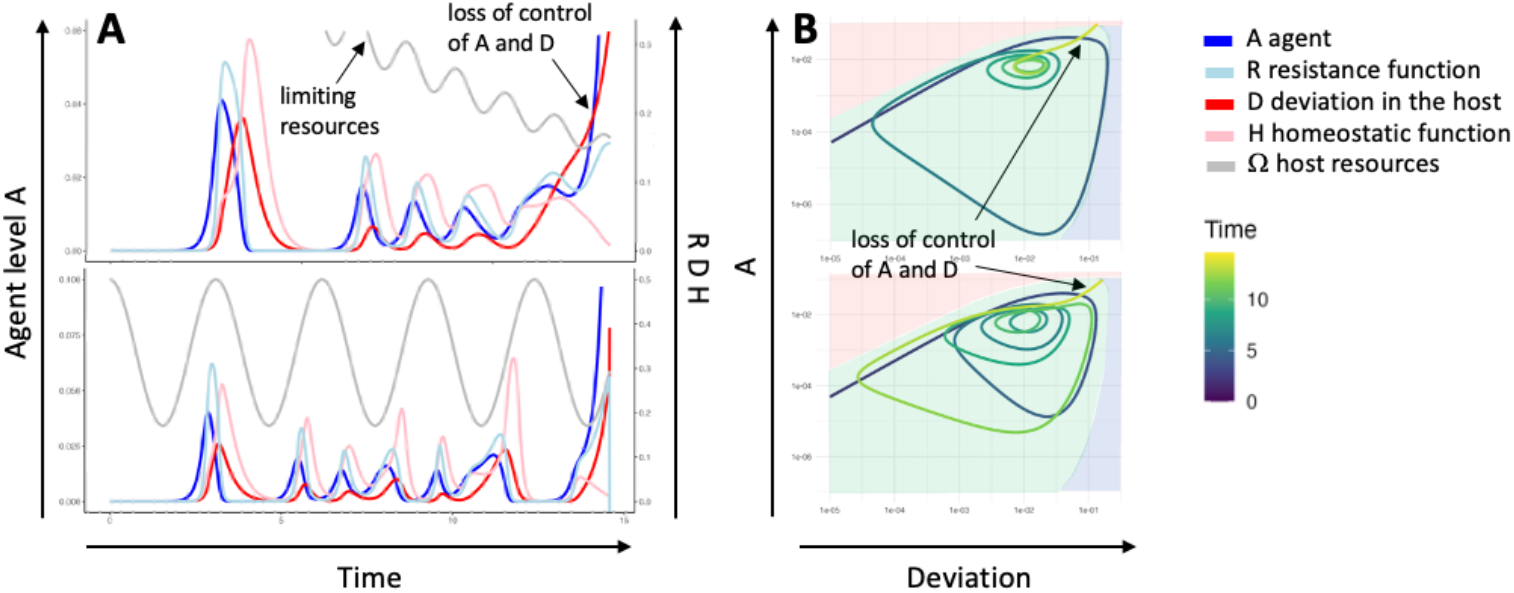
The loss of symbiosis by limiting the host’s resources. The host-microbe symbiosis can be lost by the evolution of the parameters that define the symbiosis, such as the host’s resources. Here the host’s resources fluctuate, for example as a consequence of the circadian rhythm. **A**) In the upper panel, the level of resources decreases with time, as a consequence, for example, of pathology or aging, eventually leading to a loss of control of the agent and of the deviation to the host, causing death of the host. In the lower panel, the highly fluctuating resources induce an unstable symbiosis that eventually leads to the host’s death. Parameter values are as in Figure 1, except for ρ = 10, and the resources parameters ah=4.84, bh=10, ch=0.147, dh=2.1 (upper panel), and ah=2.03, bh=2.04, ch=0, dh=0 (lower panel). **B**) The corresponding phase diagrams.

Nonetheless, here again, it is important to note that the symbiotic host-microbe relationship is devoid of valence (“symbiosis” is Greek for “living together”) and can diminish or increase the fitness of the host (which is not determined at this level, but rather by natural selection of the symbiotic relationship) ^48^. As an example, carriage of herpesviruses can confer increased resistance of the host to superinfections (a mutualistic symbiosis), but at the same time confer a lethal risk of infection in the event of subsequent immunosuppression (a parasitic symbiosis) ^21,22^. It has also to be noted that symbiosis comes with a cost to the host, as resources have to be allocated to the homeostatic and resistance functions to maintain the host-microbe equilibrium.

An important consequence of symbiosis is “colonization resistance” ^49^. As deviation is not zero during symbiosis, colonization by a second microbe is met early with a more robust resistance function (fueled by deviation), complicating the development of symbiosis of the host with new microbes. Thus, complex symbiosis (with more than one microbe) may only be established in three (self-sufficient) conditions. First, the primary symbiosis evokes low or no deviation by “dormant” microbes. However, the existence of genuine dormancy in time, or latency, which does not elicit any homeostatic or resistance function by the host, is difficult to substantiate ^22^. Second, different microbes colonize simultaneously, a situation that may occur shortly after birth. Third, different microbes colonize distinct host niches, eliciting compartmentalized host resistance functions. In support of this concept, bacterial symbionts are associated with a characteristic nutritional or spatial niche ^49-52^. For example, in the mammalian intestine, *Akkermansia mucinophila* resides in the intestinal mucus ^53^, Segmented Filamentous bacteria (SFB) reside in close proximity to intestinal epithelial cells ^54^ and *Acinetobacter sp*. reside in colonic crypts ^55^, and induce distinct types of immune responses. Conversely, compartmentalization implies that symbiosis in one niche does not guarantee symbiosis in another niche, thereby reinforcing the agent’s niche identity.

## FROM SYMBIOSIS TO SELF

What happens to the host once the microbe-host relationship is symbiotic? Does the host still react to symbiotic microbes? Once the microbe is stably integrated into the host, can it be considered part of the host’s (intuitively defined) self?

Let us consider two examples of self-components, e.g. mitochondria ^56-58^ and segments of our genome that code for endogenous retroviruses (ERVs) ^59-61^, which we now understand to be of microbial origin. The state of mitochondria in cells is tightly monitored by PRRs that are activated by the presence of mitochondrial DNA and RNA in endosomes (TLR9) and cytoplasm (cGAS-STING, RIG-1, MDA5 and the NLRP3 inflammasome), and induce the production of type I interferons, IL-1β, autophagy (in this case mitophagy) ^62,63,64^, pyroptosis (a form of programmed cells death) and the recruitment of myeloid cells to dispose of damaged cells. These PRRs are also involved in the detection of ERV activation (i.e. transcription and reverse-transcription), if these escape epigenetic silencing through DNA methylation and histone modification (involving TRIM28 and SETDB1 ^65,66^). Strikingly, the same pathways are activated by the infection of cells by bacteria or viruses, or to repress their transcriptional activity ^65,67^.

The symbiotic host-microbe relationship described in Figure 2 reflects this situation: mitochondria and ERVs are potential agents of change that are monitored by sensors of the host’s resistance function, such as PRRs. Nevertheless, if we consider mitochondria and ERVs as part of self, this situation may be regarded as a homeostate. The microbe is no more an agent of change, but is part of self, and the resistance function is no more reacting to an agent of change, but to parts of self and deviation from homeostasis: it has become a homeostatic function (Figure 5). Thus, perturbation of mitochondria homeostasis by infection, toxins or nutrient stress activates the homeostatic function to restore cellular homeostasis (via for example mitophagy) or tissular homeostasis (via elimination of the deviating cell by macrophages) ^68^. The distinction between the two functions can indeed become very blurry in the context of infection, a blurriness that nourished the debate about PRRs being receptors of (non-self) microbes or receptors of modified (or endangered) self. In fact, most PRRs recognize both types of ligands, termed MAMPs or DAMPs, respectively ^37,38^. It may be proposed that some DAMPs have a symbiont origin. It is also possible that PRRs have emerged during evolution as self “quality-control” receptors, as we will discuss below in the context of ontogeny.

**FIGURE 5.**
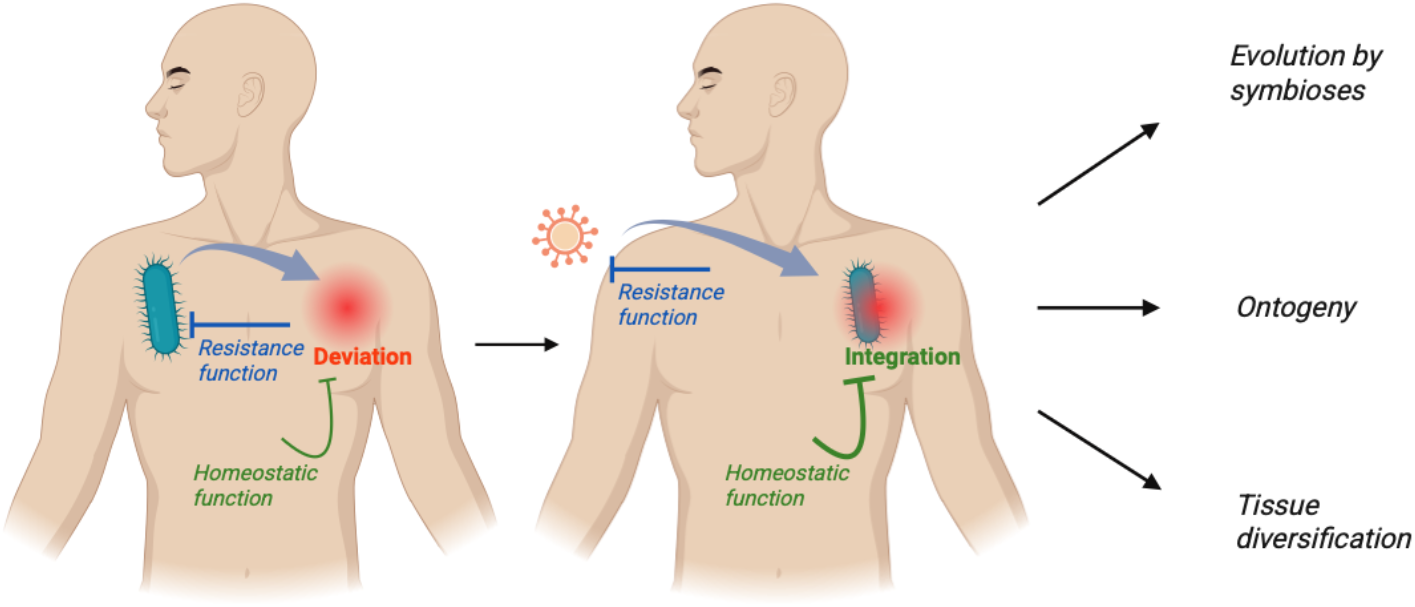
Symbiotic integration of the agent and evolution of the host. The establishment and maintenance of a symbiotic host-microbe relationship can be regarded as an integration of the microbe into “self”. In this context, the resistance function of the host against the agent becomes a homeostatic function to maintain the agent in the host. A new agent that induces deviation to the symbiotic integration will activate this homeostatic function, as well as a resistance function to the new agent. This mechanism of symbiotic integration of microbes allows for evolution of the host (examples are mitochondria and ERVs). Similar mechanisms may be at play for the evolution of complex ontogeny and tissue diversification (see text).

As discussed earlier in this paper, symbiosis does not imply a gain for the host (or the microbe) ^48^. As a matter of fact, it carries a risk for the host if the host is dependent on this symbiosis for its health and fitness, as the parameters of the symbiosis may change and lead to the loss of the symbiont (Figures 2-3, S5). For example, the loss in diversity of the intestinal microbiota is associated with host pathology in a probable circular causality of negative effects, such as, for example, the loss in producers of short chain fatty acids (SCFA) and the increase in intestinal inflammation ^69^. On the other hand, a loss of control of symbionts can lead to severe pathology ^70^, and the loss of control of endosymbionts, i.e. mitochondria ^71,72^ or ERVs ^73-75^, is associated with an array of inflammatory pathologies. Eventually, the valence of a particular symbiosis is determined by the fitness of the host by natural selection ^48^. Given the enormous number of host-microbe interactions and the diversity of symbiotic relationships they may generate, microbial symbiosis is likely to be an unavoidable and powerful, yet under-appreciated driver of evolution (as compared to genomic DNA mutations) ^76^. Importantly though, evolution through symbiosis involves modifications of the host genome by the incorporation of fragments of the microbe’s DNA ^77,78^, which facilitates vertical transmission of novel “symbiotic” traits.

## SYMBIOSIS AND ONTOGENY

Ontogeny is based on the regulated expression of genes in time and space. The product of ontogeny classically defines self ^79,80^ (even though “self” is a difficult concept to define ^81^). However, considering that stretches of our genome codes for ERVs and mitochondria, the definition of self may appear blurred from the start (of ontogeny). Furthermore, the developing fetus is characterized by the rapid generation of new molecules, cells and tissues, which must be “quality-controlled” in order to avoid pathogenic or abortive ontogeny ^82^. From the point of view of the fetal host, this process may be sensed as a succession of new (cellular) agents of change that are programmed to establish a symbiotic cell-fetus relationship, and that must be stabilized by the fetal host’s resistance and homeostatic functions to allow progress in ontogeny.

In the fetus, quality-control mechanisms rely on both (and probably complementary) cell-extrinsic and cell-intrinsic mechanisms, such as macrophages ^82^ and Toll receptors, respectively. While the functional homolog of mammalian Toll-like receptors (TLR)-type of PRRs in *Drosophila melanogaster*, Toll-1, is involved in dorsoventral patterning ^83,84^ and immunity to microbes ^85,86^, the related Toll-2, 3, 8 and 9 receptors are involved in cellular competition in wing imaginal discs, where unfit “loser” cells are eliminated by apoptosis ^87^. Similarly, the expression pattern of Toll-2, 6, 7 and 8, defines a cell surface code that is deregulated in developmentally aberrant or oncogenic cells, and modulate cellular dynamics via actomyosin and apoptosis ^39,40^. In these contexts, the Toll receptors appear to activate the host’s resistance function to the generation and positioning of a new cell type. However, in contrast to the host-microbe relationship, the resistance function is integrated in the new cell (the agent of change) itself. It is conceivable that this cell-intrinsic resistance function, performing a local quality control of the new cell type, predates the cell-extrinsic resistance function against microbes or other cells, as a way to ensure transgenerational homeostasis.

ERVs and mitochondria also play a key role in ontogeny and cell-intrinsic quality control. On the one hand, ERVs code for important elements in ontogeny, such as syncytin-1 and 2 that mediate cell-cell fusion of trophoblasts to form the syncytiotrophoblast in the placenta ^88^, HERV-K that is expressed during neuronal differentiation ^89^, and HERV-H involved in the maintenance of cell stemness ^90-93^. However, the transcription of ERVs must be tightly regulated by DNA methylation and histone modification involving the histone methyltransferase SETDB1 and the epigenetic corepressor TRIM28 ^65,66^. If deregulated, ERVs expression activates the sensors of the cellular homeostatic function discussed above (which are also sensors of the resistance function in other contexts) and induce loss of stemness, cellular differentiation or apoptosis ^93,94^. Similarly, mitochondrial activity plays a critical role in the development of cells and tissues that require a high level of energy (obtained through oxidative phosphorylation), such as neurogenesis and axonal outgrowth, cardiac tissue and myoblast differentiation, and intestinal epithelium ^95-98^. However, oxidative phosphorylation comes with the production of reactive oxygen species (ROS), which, similar to ERV products, activate the cell’s homeostatic function, induce loss of stemness, cellular differentiation or apoptosis ^97,99^. In these cases, activation of the homeostatic function leads to modification of the cellular state of homeostasis (through differentiation) or elimination of the cell if the deviation is too large.

## SYMBIOSIS AND ADAPTIVE IMMUNITY

In an insightful assay published in 2007, Margaret McFall-Ngai asks why the vast majority of animal species have evolved and thrive without the benefit of adaptive immunity ^44^. While innate immunity comes with 10-100 of PRRs (depending on the species) to recognize microbes, adaptive immunity can generate several millions of unique receptors ^100^. It is therefore widely believed that adaptive immunity provides vertebrates with a massively more robust immune system. In contrast, McFall-Ngai proposes that the vast expanse of receptors provided by adaptive immunity primarily allows vertebrates to establish more complex symbioses with microbes than invertebrates.

We have discussed earlier that complex symbioses require the colonization by different microbes of different host niches, which must elicit compartmentalized host resistance functions because of deviation to homeostasis induced by resident microbes. We also discussed that such deviation is akin to memory, encoded in the host as tissue damage, tissue memory or immunological memory, and thus, the more diverse the niches and the memory, the more discriminating the host can be in its resistance responses to microbe. As a consequence of adaptive immunity, more complex symbiotic host-microbe relationships may be established, more options may be subjected to natural selection and evolution, and more diverse cells and tissues may be generated during ontogeny.

The diversity of adaptive immunity is expressed by B cell receptors (BCR and their soluble form as antibodies) and T cells receptors (TCR), the diversity of which is generated by genetic recombination of multiple gene segments ^100^. The stochasticity of the process allows for the generation of a vast repertoire of receptors, the utility of which is selected by the recognition of a ligand and expansion of the cell producing it ^101^. As a consequence, the large intestinal microbiota induces a highly diverse repertoire of antibodies and TCRs capable of strain discrimination and control through activation of the host’s resistance function ^102-104^. Perturbation of this control leads to modifications of the microbiota with significant consequences for the host ^69,105,106^.

Interestingly, particular antibodies and TCRs have been selected through evolution to escape the risk of their loss by stochasticity, and thus, similar to PRRs, are conserved between individuals and species ^107^. For example, so-called natural (B-1) B cells produces antibodies to molecules expressed by a wide array of bacteria, such as lipopolysaccharides (LPS), peptidoglycan and phosphorylcholines, invariant natural killer T (NKT) cells recognize bacterial glycolipids ^108^, mucosa-associated invariant T (MAIT) cells recognize metabolites from the vitamin B2 (riboflavin) biosynthesis pathway produced by bacteria and fungi ^109^, and a subset of CD8^+^ T cells recognize formylated peptides characteristically produced by bacteria ^110^. These receptors may have been positively selected by evolution for their homeostatic function in maintaining specific and symbiotic (mutualistic or parasitic) host-microbe relationships.

CD8^+^ T cells that recognize formylated peptides derived from bacteria also recognize formylated peptides generated by mitochondria, and thereby induce homeostatic functions upon tissue injury ^111^. Natural B cells also produce antibodies to self-antigens involved in the clearance of apoptotic cells ^112,113^, and subsets of Tαβ and Tγδ cells recognize tissue-specific self-antigens, such as Skint1 on skin cells ^114^, presumably associated with cellular stress. These receptors may thus contribute to the diversification of the sensors required to detect the growingly diverse deviations from homeostasis in growingly complex animals.

Finally, in the thymus, self-reactive immature T cells differentiate into so-called regulatory T cells (Tregs), adding a large pool of T cells and receptors that can contribute to the host’s homeostatic function ^115-117^. Tregs can also be generated from T cells that react against microbial symbionts ^118,119^. The importance in the homeostatic function of this pool of anti-self (or anti-symbiont) T cells is underscored by the lethal multi-organ inflammation that develops in human and mouse carrying deleterious mutations in *Foxp3*, coding for a transcription factor necessary in the generation of Tregs ^120-122^. Even though this effect is generally interpretated as a failure of Tregs to inhibit autoimmunity (by auto-reactive effector T cells and B cells) or inflammatory disease of mucosal tissues (such as inflammatory bowel disease, IBD, involving symbiont-reactive effector T cells), it may also be viewed as a failure of the homeostatic function to maintain cell, tissue and organ homeostasis ^116,117^, thereby fueling the resistance function against colonizing microbes (in mucosal or internal tissues), or modifying the conditions that allowed for symbiosis with microbes, mitochondria, ERVs, or other self-components, and turning homeostatic functions “back” into resistance functions ^71-75^. Even though the two interpretations lead to the same effects, the solutions they propose for cure are different. Following the “autoreactivity” interpretation, the generation of auto-reactive effector cells must be inhibited or redirected into Tregs. In contrast, following the “homeostasis” interpretation, it is the agent of change (microbe, symbiont), and the deviation it induces to the host (such as tissue damage), that must be controlled. In this view, blocking the resistance function (autoimmune T cells) in favor of the homeostatic function (Tregs) may lead to a loss of control of symbionts and own cells, and induce an increase in inflammatory pathology and cancer.

## DISCUSSION

The formal model of host-microbe relationship presented here is constructed on the principle that homeostasis is a core attribute of life. To maintain homeostasis in the face of environmental variations, microbes and internal dynamics, we propose that the host needs both a homeostatic function to counter change induced by an agent ^26-28^, and a resistance function to counter the agent of change directly. We show that these two functions are sufficient to generate a symbiotic host-microbe relationship, by avoiding the host’s lethal deviation from homeostasis induced by the agent, or elimination of the agent from the host. Furthermore, as symbiosis allows for the (beneficial or detrimental) “integration” of the agent into the host, we propose that the same mechanisms offer options for evolution by natural selection of the host-microbe symbiosis, as well as for the management of growingly complex ontogeny in metazoans.

We show here that symbiosis is an emergent property of the host-microbe relationship, which does not require the microbe or the host to express particular “traits of symbiosis”. Emergence relates to properties observed in a system that are not present in individual parts of the system, and that cannot be explained solely from these individual parts ^123-126^. Contrary to this principle, a more reductionist approach of microbiology and immunology has dominated the study of host-microbe relationships, on the assumption that microbes are intrinsically pathogens or symbionts to the host, or pathobionts if they reveal their pathogenic nature only under certain conditions ^9-11^, and that the immune system is equipped with cells, such as Tregs, that provide symbiont-specific tolerance ^118,119,127^. We rather conclude that symbiosis is a natural consequence of host-microbe relationships that follow the rule of homeostasis, a key emergent property of living organisms ^26,27^.

Nonetheless, we opened this paper with the frequently asked question of “how the immune system discriminates a pathogen from a symbiont”, and challenged the notion that this was a good question. Given that a particular microbe can be mutualistic or parasitic at different times or niches during the host-microbe relationship ^21-24^ we and others proposed that a microbe is not intrinsically pathogenic or symbiotic, but rather that the host-microbe relationship is pathogenic or symbiotic ^18,19,25^. This view is, in some ways, similar to that of the “danger theory” ^128,129^ stating that a microbe activates the immune system through the danger it represents to the host, expressed at the molecular levels by proxys of danger or released by damaged tissues, cells or molecules as DAMPs ^37,38,129,130^. We incorporate this concept in our model as a broader (and neutral) notion of deviation to the host, which is required to fuel the resistance function of the host to the microbial agent of change.

More specifically however, in the host-microbe relationship, the danger theory proposes that DAMPs signal pathogenicity. Pathogens would generate DAMPs, because, for example, of penetration into “forbidden” compartments, such as the cytoplasm ^13^, and the mode of eventual cell death they induce ^131^. Symbionts, in contrast, would reduce the generation of DAMPs by promoting the host’s homeostatic (in this case, repair) function ^8,73,111,132,133^. Nevertheless, cytoplasmic residence of microbes is not a rare occurrence in eukaryote cells ^78,134^, and damage is not specific to pathogens as symbionts also induce damage ^135-138^. Mitochondria, a descendent of α*-Proteobacteria*, is maybe the most striking example of such stable residence ^58^, which can generate substantial damage by the production of ROS, nonetheless “tolerated” to some level by the host’s homeostatic function ^28^. We therefore argue that damage is a common occurrence in living systems that cannot determine the nature of a microbe, and we thus include the notion of damage, in our model, into the broader and neutral notion of deviation to the host.

In a somewhat circular (tautological) argument ^139^, DAMPs are defined by the “pathological” inflammation (and clinical pathology) they induce ^37,38,129^, whereas “physiological” or “homeostatic” inflammation is involved in the maintenance of homeostasis (such as repair responses) ^4,7,8^. However, while the pathological state of a tissue is relatively straightforward to measure, the “pathological” or “physiological” nature of inflammation at the cellular and molecular levels is not. For example, robust “homeostatic” type 3 responses are necessary in the intestine to maintain an equilibrium with the large intestinal microbiota ^1-6,140,141^, and this type of responses is similar to pathological type 3 responses developing in the context of autoimmunity ^142-144^. This is valid for all types of immune responses, which are involved in the maintenance of symbiosis with different types of microbes, as well as in different types of pathological inflammation ^24^. Attempts are made to differentiate, for example, physiological versus pathological type 3 immune responses in the intestine, at the cellular and molecular levels ^145-150^, but fail to identify a “red line” of mediators specifying pathological inflammation. In our model, we define a resistance function against the microbe, and a homeostatic response against the damage induced by the microbe, also termed tolerance to the microbe ^43^. However, we refrain from associating the former to “pathological” and the latter to “physiological” or “homeostatic”, as we predict that the level of resistance function can be low (and apparently physiological) to eliminate a microbe (Figure 1), or high (apparently pathological) to maintain symbiosis (Figure 2).

Another problematic issue is the definition of symbiont. On the basis of the original Greek definition of symbiosis, i.e. “living together”, a symbiont can be pathogenic. Indeed, if “living together” in the host-microbe relationship means a prolonged host-microbe interaction, then notorious pathogens, such as HIV and *Mycobacterium tuberculosis*, are efficient symbionts, or “pathobionts” ^151^. Furthermore, if we use the more common notion of symbiosis as a mutualistic host-microbe interaction, then we next have to define mutualism. For example, symbionts in our intestine provide benefits to the host through food digestion and maintenance of the gut barrier, but can become pathogenic if food and the host’s condition change ^152,153^. In another example, herpesviruses may protect from superinfection or act as pathogens by overwhelming the host, depending on the state of the host ^21-23^. Of note, it is more straightforward to define a pathogen as a cause of disease, using the Koch postulates to demonstrate its pathogenic nature, than to identify its mutualist nature when the host is healthy (and nothing happens to the host at the clinical level). But while a clinician identifies a pathogen because it makes their patients sick, how can the host, and its immune system, know that a microbe, such as HIV and *Mycobacterium tuberculosis*, will eventually (years later) kill it? We thus propose that the host cannot make this distinction, but merely measures the level of deviation to homeostasis induced by the microbe, and counters such deviation by engaging its homeostatic and resistance functions ^28^. Both notions of “deviation” (such as damage) and “agent of deviation” (here the microbe) are neutral, because the valence of deviation is contextual, and eventually selected by the environment of the individual, as are all individual traits.

Our model shows that the limit between symbiosis or the eventual loss of microbe or death of the host is an infinitesimal small line, best described by chaos theory ^45-47^, where a small change in parameters has a dramatic effect on the outcome in the host-microbe relationship. The trajectory in time of this relationship can be described by a phase diagram (Figures 2-4) as a function of the level of agent (A, or microbe) in the host, and the level of the deviation (D) to homeostasis it induces in the host. A useful development of this view would be to define biologically and clinically relevant measures of A and D, as well of the attraction basin to symbiosis, to predict the evolution of the host-microbe relationship. Quantification of a specific agent in the host seems possible, at least in defined (cellular, tissular, organ) niches. Quantification of deviation may also be possible, once the relevant deviation induced by the agent is characterized. Definition of the attraction basin may be more complex, as it is a function of many parameters (in this model), and thus, may have to be defined empirically.

We have developed a general model of homeostasis and applied it to the particular case of host-microbe interactions, without any notion of “good” or “bad”, but with the central assumption that homeostasis is fundamental to the host. The model also allows to make a number of propositions on the host-microbe relationship and immunity (Box 1), and to examine specific contexts of pathogenesis generated by modification of this relationship (Figure 5). It may be interesting to apply this model to other types of host-agent relationships, where the agent of change is, for example, an allergen, a psychological stressor or an organism kin to the host. In the case the agent of change is an allergen, it will fundamentally differ from a microbe as being non-replicative. In the case the agent of change is a psychological stressor, the agent will induce deviations of the mental state, to which the homeostatic function responds by toning down the effect of that stressor (by the action, for example, of external or internal cannabinoids), while the resistance function will respond by attempting to counter the stressor. In these cases, formalization of the agent’s behavior, and of the host’s homeostatic and resistance functions to it, will require a new set of equations describing A, H and R.

It will be fascinating to explore the host-agent relationship in the case the agent is kin, such as another person, and thus establish the conditions of symbiosis that avoid elimination of the agent (the other person), the cost of maintaining the symbiosis, and the risks brought by this symbiosis if the parameters defining the agent and the host are altered. Of note, the model presented here, as applied to the host-microbe relationship, is host-centric, as we consider only the host’s homeostasis. A symmetrical model can be developed to take in account the microbe’s own homeostasis, a symmetry that would be necessary to explore social relationships.

## ACKNOWLEDGEMENTS

I would like to thank Thomas Pradeu for his broad knowledge of the historical and philosophical aspects of immunology, and the comprehensive discussions we have on symbiosis, Olivier Mirabeau for his help on coding and data visualization, Laure Bally-Cuif for her decisive insights into ontogeny and the role of Toll receptors in quality control, Romain Levayer for his generous insights in cellular interactions during ontogeny, Gabriel Lepousez for his remarks on systems behavior and phase diagrams, Marc Daëron for our initial discussions on symbiosis, Philippe Kourilsky for his interest in more general views of immunity, Philippe Sansonetti for his defense of microbes, Kama Atretkhany for her remarks on energy and host resources, Paulo Vieira for his interest in new views of immunity, Yasmine Belkaid for our shared interest in defining immunity and symbiosis, Hannah Kaminsky for her work and insights on symbiosis and repair, Fridolin Gross for his insights in mathematical formulation and discussions on the Lotka-Volterra equations, Samir Ali Moussa for our discussions of the model, and the Microenvironment & Immunity lab for feedbacks on this work.

**Box 1: Propositions generated by the model**

## Microbial agents

### 1. Low impact microbes can be lethal

Since the impact of the microbe on the host fuels host resistance, low impact (defined in this model as inducing limited deviation to the host) microbes may expand excessively and, eventually, induce more extensive damage and strong immune responses that are immunopathogenic. Importantly, the impact of a microbe on the host is not an intrinsic property of the microbe, but a property of the microbe-host relationship. Example: the host-dependent effect of SARS-CoV-2 ^41,42^.

## The homeostatic function

### 2. Strong homeostatic (regeneration and repair) functions can be lethal

Even though it is intuitively beneficial to reduce as fast and as much as possible the damage generated by a microbe, too much repair may lead to excessive tissue tolerance to the microbe, and, as a consequence, to a potentially pathogenic loss of microbe control.

### 3. Mid-level homeostatic (regeneration and repair) functions are key to symbiosis

Intermediate levels of homeostatic functions prevent, on the one hand, too high a tissue deviation (damage) that fuels resistance to, and eventually the elimination of, the microbial agent, while on the other hand, too fast a return to homeostasis leads to a loss of microbe control. Thus, within these two extremes, the homeostatic function allows for a state of equilibrium (allostate) between microbe and host, defined as symbiosis.

## The resistance function

### 4. The resistance function is required for immunity, symbiosis and self

The resistance function, by definition, targets the (microbial) agent, and its level is a function of the number of microbes. However, resistance is not immunity: the resistance function is key for microbe elimination, but also for microbial control during symbiosis. Once a stable equilibrium (symbiosis) is established between the microbe and the host, the microbe appears to become functionally self, and the resistance function becomes a homeostatic function, as it is required to maintain this new state.

## Symbiosis

### 5. Symbiosis is an allostate resulting from the homeostatic and resistance functions

Paradoxically, the homeostatic function that opposes change, and the resistance function that opposes the microbial agent of change, are key to establish the microbe-host symbiosis. Importantly however, a microbe-host symbiosis has no intrinsic positive or negative value, as it is merely the consequence of the interaction between a microbe and its host.

### 6. Symbiosis is a source of pathology

Changes to the microbe or the host, through mutations, modifications of diet, niche or energy levels, or medication, may destabilize the symbiotic allostate and lead to the loss of the microbe, or lead to pathogenic expansion of the microbe (Figure S5).

### 7. Symbiotic allostates are drivers of evolution

The establishment of successive symbiotic allostates is a powerful driver of evolution through the incorporation into the host of microbial agents (such as mitochondria and retroviruses).

### 8. Ontogeny is a rapid succession of symbiotic allostates

The succession of multiple allostates may also describe the ontogenic process that continuously incorporates new components into the developing host. The homeostatic and resistance functions are key in this process to stabilize these (symbiotic) additions, preventing their loss or lethal overgrow.

## Adaptive immunity

### 9. Adaptive immunity serves immunity, symbiosis and ontogeny

Adaptive immunity generates a very large repertoire of receptors that may not only serve the resistance function to invasive microbes (immunity) but also the homeostatic function (regeneration and repair). It thereby stabilizes complex (multiple) symbioses with microbes, as well as the high tissue diversity in vertebrate hosts ^44^.

### 10. Tregs contribute fundamentally to the homeostatic function

Although classically characterized as opposing the resistance (pro-inflammatory) function, the fundamental role of Tregs may be as contributors to the homeostatic (regeneration and repair) function ^116,117^. The net (clinical) effect of Tregs on the host is nevertheless similar (preventing autoimmunity), as in this view, Tregs decrease the adjuvant effect of tissue damage on the resistance function.

## SUPPLEMENTARY MATERIALS

**Figure S1. Limitations in time and energy (Ψ).**

To include a logistic evolution in time of the homeostatic (H) and resistance (R) functions, we introduce the logistic function S:

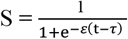 in their description, where t is the time, τ the time delay to adjust S ∼ 0 at t = 0, and ε the rate of acceleration. For the sake of simplicity, the same values for τ and ε have been used here to describe H and R. Different values of τ and ε for H and R lead to quantitative but not qualitative differences to the model.

To express a limitation in the host’s energy available for the homeostatic (H) and resistance (R) functions, we introduce the negative feedback function Ω:

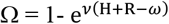 in their description, where *ω* is the physiological limit of resources, which we set as ω = 1, and *ν* the rate of the negative feedback. In the main text, Ψ = SΩ.

The value of ω may change as a consequence of the host’s circadian rhythm, physiological stress and age, during which energy is allocated to other physiological functions, or lost. To this end, in the R code reported in figure S4, we have introduced:

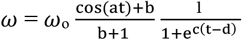 in the description of Ω, where a is the frequency of oscillation in energy availability, and b the tolerance to oscillations, in order to create a context of rhythmic availability of energy to the H and R functions. Furthermore, c is the rate of decrease, and d the timing of decrease, in order to create a context of decreasing availability of energy, for example with age.

**Figure S2. System of equations describing deviation from homeostasis in the host-microbe relationship**

We obtain a system of 4 equations that describe the evolution in time of the agent (A) in the host, its impact on the host’s deviation (D) from homeostasis, of the host’s homeostatic function (H) and of the resistance function (R) against the agent:

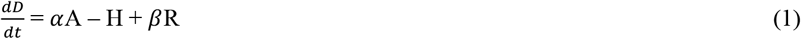

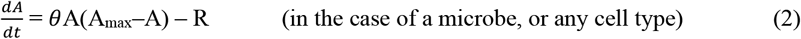

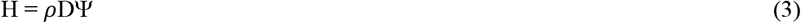

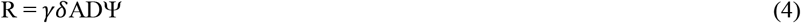

with:

D the deviation from homeostasis

*α* the rate of damage induced by the agent A the amount of agent

H the level of homeostatic function

R the level of resistance function targeting A

*β* the deviation induced by the resistance function (R)

*θ* the intrinsic growth rate of A

A_max_ the maximal amount of A in the host (the niche)

*ρ* the efficacy of the homeostatic force

*γ* the reactivity of the response to A

*δ* the adjuvant effect of the deviation on the resistance function (R)

Ψ the corrective function for the limitation in time and resources (Figure S1) This system of equations is solved by the deSolve package on R (Figure S3).

Importantly, Ψ, and more specifically Ω, is a problem when attempting to resolve the system of equations (1-4) as Ω is determined by the values of H and R (Ω = 1-e^*ν*(H+R-*ω*)^)and the values of both H and R depend on Ω (equations 3 and 4). Thus, we first have to find the descriptions for

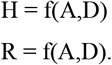

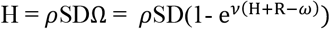 which leads to : 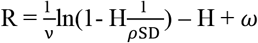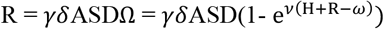 which leads to: 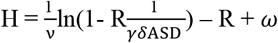 Combining the two developments, we obtain:

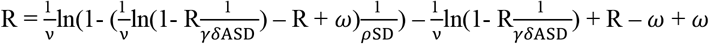

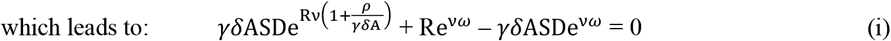

and 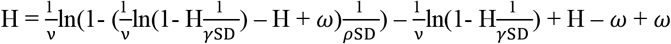

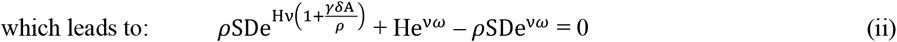

Both equations (i) and (ii) are of the form ae^bx^+cx + d = 0 δ

The solution to such equations is obtained with Lambert’s W(w) function, where:

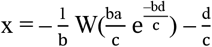

For (i) and x = R: a = *γδ*ASD, 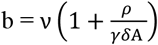, c = e^*νω*.^, d = – *γδ*ASDe^*νω*.^

For (ii) and x = H: a = *ρ*SD, 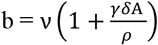, c = e^*νω*.^, d = – *ρ*SDe^*νω*.^

Finally, the quantitative, but not the qualitative behavior of this system of equations does not change if, for example, H and R are non-linear functions of D and A:

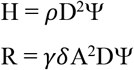

The reader can assess this by modifying the R code modeling the system of equation 1-4 (Figure S3).

We provide here a table summarizing the behavior of A, D, H and R, as well as of the reaction time, in response to an increase in each individual parameter.

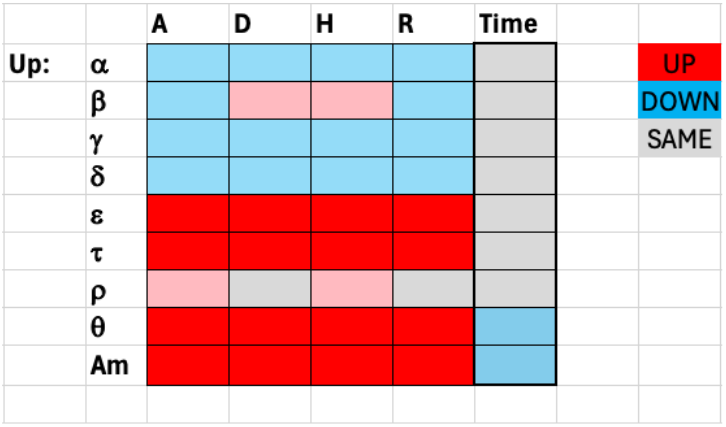

**Figure S3a. R code to visualize the evolution with time of A, D, H, R**

~~~
install.packages(“devtools”)
install.packages(“shiny”)
install.packages(“shinySIR”)
install.packages(“deSolve”)
install.packages(“tidyverse”)
install.packages(“lamW”)
library(devtools)
library(shiny)
library(shinySIR)
library(deSolve)
library(tidyverse)
library(lamW)
# MODEL FUNCTION
dar.model <-function(t, state, parameters) {
 with(as.list(c(state, parameters)), {
  # Avoid underflow when A is very small
  A_eff <-max(A, 1e-12)
  # Common terms
  S <-1 / (1 + exp(-ep * (t - ta)))
  om_t <-om * ((cos(ah * t) + bh) / (bh + 1)) / (1 + exp(ch * (t - dh)))
  # Core terms
  a1 <-rh * S * D
  a2 <-ga * de * A_eff * S * D
  b1 <-nu * (1 + (ga * de * A_eff / rh))
  b2 <-nu * (1 + (rh / (ga * de * A_eff)))
  c <-exp(nu * om_t)
  d1 <- -rh * S * D * c
  d2 <- -ga * de * A_eff * S * D * c
  # Responses
  H <- -1/b1 * lambertW0(b1 * a1 / c * exp(-b1 * d1 / c)) - d1 / c
  R <- -1/b2 * lambertW0(b2 * a2 / c * exp(-b2 * d2 / c)) - d2 / c
  # Differential equations
  dD <-al * A_eff + be * ga * de * S * D * A_eff * (1 - exp(nu * (R + H - om_t))) –
  rh * S * D * (1 - exp(nu * (R + H - om_t)))
  dA <-ifelse(A > 1e-12,
                 th * A * (Am - A) - ga * de * S * D * A * (1 - exp(nu * (R + H - om_t))), 0)
  list(c(dD, dA))
 })
}
# SHINY APP
ui <-fluidPage(
 titlePanel(“Parameter Exploration”),
 sidebarLayout( 
  sidebarPanel(
   sliderInput(“ti”, “Time”, min = 0, max = 100, value = 7, step = 0.01),
  sliderInput(“al”, “Agent impact (alpha)”, min = 0, max = 5, value = 1.5, step = 0.01),
  sliderInput(“be”, “R impact (beta)”, min = 0, max = 10, value = 1.4, step = 0.01),
  sliderInput(“ga”, “R reactivity (gamma)”, min = 0, max = 100, value = 25, step = 1),
  sliderInput(“de”, “Adjuvant D (delta)”, min = 0, max = 100, value = 35, step = 0.01),
  sliderInput(“ep”, “Logistic speed (epsilon)”, min = 0, max = 10, value = 0.6, step = 0.01),
  sliderInput(“ta”, “Logistic time delay (tau)”, min = 0, max = 20, value = 5, step = 0.01),
  sliderInput(“rh”, “P reactivity (rho)”, min = 0, max = 30, value = 2, step = 0.01),
  sliderInput(“th”, “Agent growth potential (theta)”, min = 0, max = 10, value = 3.5, step = 0.01),
  sliderInput(“Am”, “Agent max (Am)”, min = 0, max = 10, value = 1.8, step = 0.01),
  sliderInput(“om”, “Energy limit (omega)”, min = 0, max = 1, value = 1, step = 0.01),
  sliderInput(“nu”, “Energy limit tolerance (nu)”, min = 0, max = 20, value = 10, step = 0.01),
  sliderInput(“ah”, “Frequency E oscillation (ah)”, min = 0, max = 5, value = 0, step = 0.01),
  sliderInput(“bh”, “Tolerance E oscillation (bh)”, min = 1, max = 10, value = 10, step = 0.01),
  sliderInput(“ch”, “Slope E decrease (ch)”, min = 0, max = 2, value = 0, step = 0.001),
  sliderInput(“dh”, “Timing E decrease (dh)”, min = 0, max = 20, value = 0, step = 0.01)
  ),
  mainPanel(
   plotOutput(“plot1”),
   plotOutput(“plot2”),
   plotOutput(“plot3”),
   plotOutput(“plot4”)
  )
 )
)
# SERVER FUNCTION
server <-function(input, output) {
 output$plot1 <-output$plot2 <-output$plot3 <-output$plot4 <-renderPlot({
  parms <-c(ti = input$ti, al = input$al, be = input$be, ga = input$ga, de = input$de, ep = input$ep, ta = input$ta,
                rh = input$rh, th = input$th, Am = input$Am, om = input$om, nu = input$nu,
                ah = input$ah, bh = input$bh, ch = input$ch, dh = input$dh)
   times <-seq(from = 0, to = input$ti, by = 0.01)
   xstart <-c(D = 0, A = 1e-9)
   ode(func = dar.model, y = xstart, times = times, parms = parms) %>%
    as.data.frame() -> out
   for (k in 1:length(names(parms))) {
    assign(names(parms)[k], as.vector(parms[k]))
   }
   S <-1 / (1 + exp(-ep * (out$time - ta)))
   om_t <-om * ((cos(ah * out$time) + bh) / (bh + 1)) / (1 + exp(ch * (out$time - dh)))
   A_eff <-pmax(out$A, 1e-12)
   a1 <-rh * S * out$D
   a2 <-ga * de * A_eff * S * out$D
   b1 <-nu * (1 + (ga * de * A_eff / rh))
   b2 <-nu * (1 + (rh / (ga * de * A_eff)))
   c <-exp(nu * om_t)
   d1 <- -rh * S * out$D * c
   d2 <- -ga * de * A_eff * S * out$D * c
   H <- -1 / b1 * lambertW0(b1 * a1 / c * exp(-b1 * d1 / c)) - d1 / c
   R <- -1 / b2 * lambertW0(b2 * a2 / c * exp(-b2 * d2 / c)) - d2 / c
   # Combine data
   out <-cbind(out, R, H, om_t)
   # Define color palette
   pal <-c(“blue”, “red”, “pink”,”grey”, “lightblue”)
   out_long <-out %>%
     gather(variable, value, -time) %>%
     mutate(group = ifelse(variable == “A”, “A”, “others”))
   # - -- - MANUAL LIMITS - -- -
   ymin_left <-0 # for A
   ymax_left <-0.1
   ymin_right <-0    # for D, H, R
   ymax_right <-0.5
   # Scaling factor to map right axis values into left axis space
   scale_factor <-(ymax_left - ymin_left) / (ymax_right - ymin_right)
   ggplot(out_long, aes(x = time)) +
    geom_line(data = subset(out_long, group == “A”),
        aes(y = value, color = variable), size = 2) +
    geom_line(data = subset(out_long, group == “others”),
        aes(y = value * scale_factor, color = variable), size = 2) +
    scale_y_continuous(
     name = “A”,
     limits = c(ymin_left, ymax_left),
     sec.axis = sec_axis(∼./scale_factor,
                   name = “D, H, R”,
                   breaks = seq(ymin_right, ymax_right, by = 0.1))
     ) +
     scale_color_manual(values = pal) +
     labs(x = “Time”, color = “Variable”) +
     theme_classic(base_size = 14) +
     theme(
      legend.position = “right”,
      legend.title = element_text(size = 12), 
      legend.text = element_text(size = 12),
      axis.title = element_text(size = 14),
      axis.text = element_text(size = 12)
     )
  })
}
shinyApp(ui=ui, server=server)
~~~

**Figure S3b. R code for phase diagrams**

~~~
 install.packages(“devtools”)
 install.packages(“shiny”)
 install.packages(“shinySIR”)
 install.packages(“deSolve”)
 install.packages(“tidyverse”)
 install.packages(“lamW”)
 library(devtools)
 library(shiny)
 library(shinySIR)
 library(deSolve)
 library(tidyverse)
 library(lamW)
 library(tidyr)
 library(ggplot2)
 # MODEL FUNCTION
 dar.model <-function(t, state, parameters) {
  with(as.list(c(state, parameters)), { 
   # Avoid underflow when A is very small
   A_eff <-max(A, 1e-12)
   # Common terms
   S <-1 / (1 + exp(-ep * (t - ta)))
   om_t <-om * ((cos(ah * t) + bh) / (bh + 1)) / (1 + exp(ch * (t - dh)))
   # Core terms
   a1 <-rh * S * D
   a2 <-ga * de * A_eff * S * D
   b1 <-nu * (1 + (ga * de * A_eff / rh))
   b2 <-nu * (1 + (rh / (ga * de * A_eff)))
   c <-exp(nu * om_t)
   d1 <- -rh * S * D * c
   d2 <- -ga * de * A_eff * S * D * c
   # Responses
   H <- -1/b1 * lambertW0(b1 * a1 / c * exp(-b1 * d1 / c)) - d1 / c
   R <- -1/b2 * lambertW0(b2 * a2 / c * exp(-b2 * d2 / c)) - d2 / c
   # Differential equations
   dD <-al * A_eff + be * ga * de * S * D * A_eff * (1 - exp(nu * (R + H - om_t))) - rh * S * D * (1 - exp(nu * (R + H - om_t)))
   dA <-ifelse(A > 1e-12,
                 th * A * (Am - A) - ga * de * S * D * A * (1 - exp(nu * (R + H - om_t))), 0)
   list(c(dD, dA))
  })
 }
 # SHINY APP
 ui <-fluidPage( 
  titlePanel(“Parameter Exploration”),
  sidebarLayout(
   sidebarPanel(
   sliderInput(“ti”, “Time”, min = 0, max = 50, value = 15, step = 0.01),
   sliderInput(“al”, “Agent impact (alpha)”, min = 0, max = 5, value = 1.5, step = 0.01),
   sliderInput(“be”, “R impact (beta)”, min = 0, max = 10, value = 1.4, step = 0.01),
   sliderInput(“ga”, “R reactivity (gamma)”, min = 0, max = 100, value = 25, step = 1),
   sliderInput(“de”, “Adjuvant D (delta)”, min = 0, max = 100, value = 35, step = 0.01),
   sliderInput(“ep”, “Logistic speed (epsilon)”, min = 0, max = 10, value = 0.6, step = 0.01),
   sliderInput(“ta”, “Logistic time delay (tau)”, min = 0, max = 20, value = 5, step = 0.01),
   sliderInput(“rh”, “P reactivity (rho)”, min = 0, max = 25, value = 3, step = 0.01),
   sliderInput(“th”, “Agent growth potential (theta)”, min = 0, max = 10, value = 3.5, step = 0.01),
   sliderInput(“Am”, “Agent max (Am)”, min = 0, max = 10, value = 1.8, step = 0.01),
   sliderInput(“om”, “Energy limit (omega)”, min = 0, max = 1, value = 1, step = 0.01),
   sliderInput(“nu”, “Energy limit tolerance (nu)”, min = 0, max = 20, value = 10, step = 0.01),
   sliderInput(“ah”, “Frequency E oscillation (ah)”, min = 0, max = 5, value = 0, step = 0.01), 
   sliderInput(“bh”, “Tolerance E oscillation (bh)”, min = 1, max = 10, value = 10, step = 0.01),
   sliderInput(“ch”, “Slope E decrease (ch)”, min = 0, max = 2, value = 0, step = 0.001),
   sliderInput(“dh”, “Timing E decrease (dh)”, min = 0, max = 20, value = 0, step = 0.01)
 ),
 mainPanel (
  plotOutput(“plot1”),
  plotOutput(“plot2”),
  plotOutput(“plot3”),
  plotOutput(“plot4”)
  )
 )
)
# SERVER FUNCTION
server <-function(input, output) {
 output$plot1 <-output$plot2 <-output$plot3 <-output$plot4 <-renderPlot({
 rh_values <-input$rh
 all_results <-data.frame()
 for (rh_val in rh_values) {
 parms <-c(
  ti = input$ti, al = input$al, be = input$be, ga = input$ga, de = input$de, ep = input$ep, ta = input$ta,
  rh = rh_val, th = input$th, Am = input$Am, om = input$om, nu = input$nu, ah = input$ah, bh = input$bh,
  ch = input$ch, dh = input$dh
 )
  times <-seq(from = 0, to = input$ti, by = 0.01)
  xstart <-c(D = 0, A = 1e-9)
 ode(func = dar.model, y = xstart, times = times, parms = parms) %>%
  as.data.frame() -> out
 S <-1 / (1 + exp(-parms[“ep”] * (out$time - parms[“ta”])))
 om_t <-parms[“om”] * ((cos(parms[“ah”] * out$time) + parms[“bh”]) / (parms[“bh”] + 1)) / (1 + exp(parms[“ch”] * (out$time - parms[“dh”])))
  A_eff <-pmax(out$A, 1e-12)
  a1 <-parms[“rh”] * S * out$D
  a2 <-parms[“ga”] * parms[“de”] * A_eff * S * out$D
  b1 <-parms[“nu”] * (1 + (parms[“ga”] * parms[“de”] * A_eff / parms[“rh”]))
  b2 <-parms[“nu”] * (1 + (parms[“rh”] / (parms[“ga”] * parms[“de”] * A_eff)))
  c <-exp(parms[“nu”] * om_t)
  d1 <- -parms[“rh”] * S * out$D * c
  d2 <- -parms[“ga”] * parms[“de”] * A_eff * S * out$D * c
  H <- -1 / b1 * lambertW0(b1 * a1 / c * exp(-b1 * d1 / c)) - d1 / c
  R <- -1 / b2 * lambertW0(b2 * a2 / c * exp(-b2 * d2 / c)) - d2 / c
  out <-cbind(out, R, H, om_t)
  out$rh_val <-rh_val # Tag each row with respective rh value
  all_results <-rbind(all_results, out)
 }
 # To avoid overplotting, optionally sample time points:
 sampled_results <-all_results %>% filter(time %% 0.01 < 0.01) # every 1 unit of time
 ggplot(sampled_results, aes(x = D, y = A, color = time, group = factor(rh_val))) + 
  geom_path(size = 2) +
  scale_x_log10(limits = c(1e-5, 0.2)) +
  scale_y_log10(limits = c(1e-12, 0.1)) +
  scale_color_viridis_c(name = “Time”) +
  labs(x = “D”, y = “A”, title = “A vs D colored by time”) +
  theme_minimal(base_size = 14)
 })
}
shinyApp(ui=ui, server=server)
~~~

**Figure S4. Conditions to establish symbiosis**

To establish symbiosis of the host with a microbe, two (obvious) conditions must be met. First, the microbe must not be eliminated, and second, the host cannot die. Mathematical treatment of these two conditions leads to two inequations.

*1. Condition to avoid the host’s elimination of the agent (lowest levels of A)*

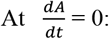

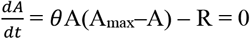 so *θ*A(Amax–A) = R or *θ*A(Amax–A) = *γδ*AΨ which leads to 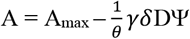

We set here a lower limit for A as A_lim_ > 10^-12^ (we stipulate that less than 10^-12^ microbes in the host would mean less than one microbe). So symbiosis can only be established if 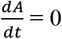 occurs before A_min_. So: 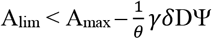 and *γδ*DΨ < *θ*(A_max_–A_lim_) As A_max_ >> A_lim_ we can write A_max_–A_lim_ ≈ A_max_ and:

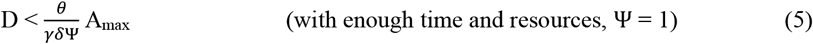

*2. Condition to avoid the agent’s killing of the host (highest levels of D)*

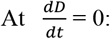

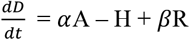 so 0 = *α*A – *ρ*DΨ + *βγδ*DAΨ which leads to 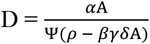

We set here the higher limit of D so that D < 1 (we stipulate that D = 1 means death of the host). So, symbiosis can only be established if 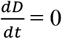 occurs before D = 1. So: 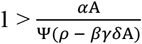 and:

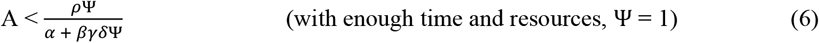

**Figure S5. From symbiosis to pathogenesis**

To establish symbiosis of the host with a microbe, two conditions must be met. First, the microbe must not be eliminated, and second, the host cannot die. Mathematical treatment of these two conditions leads to two inequations that must be satisfied to establish and maintain symbiosis (see Figure S4):

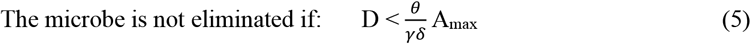

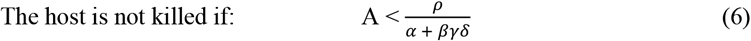

Nevertheless, symbiosis can be source of pathology if the agent’s nature or the host’s functions are altered. Five different pathogenic scenarios may unfold in the context of a host and its symbiotic microbes:

1. A first scenario derives from inequation (5) 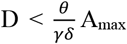 a decrease in microbial fitness (defined by q and A_max_, upon a change in the host’s diet, tissue niche or antibiotic treatment, for example) or an increase in host resistance against the microbe (defined by γ and δ, upon a loss in immunoregulation after infection, for example), may lead to the loss of the agent. With the loss of the agent, the host may lose important factors for health). Example: the loss of species and functional diversity in the microbiota of patients developing inflammatory bowel disease (IBD) ^154^.
2. The other scenarios derive from inequation (6) 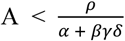, i.e. an increase in deviation D to levels damageable to the host: first, the level of the microbial agent A increases as a consequence of increased fitness upon a change in the host’s diet or niche tissue, or genetic modifications in the agent. Example: the expansion of *Candida albicans* with carbohydrate supplementation ^155^.
3. Alternatively, the impact of the agent on the host (defined by *α*) increases, again, for example, as a consequence of changes in the host’s diet or niche tissue, or genetic modifications in the agent. Example: an increase in host temperature activates the expression of virulence factors in *Shigella* ^156^.
4. The host’s resistance to the microbial agent (defined by γ and δ) increases, as does “immunopathology” (defined by *βγδ*), as a consequence of a loss in immunoregulation. Example: the loss of the immunoregulatory factor IL-10 leads to increased resistance of the host to the intestinal symbiont *Helicobacter hepaticus*, and leads to intestinal immunopathology ^157^.
5. The host’s homeostatic function (defined by *ρ*) decreases, as a consequence, for example, of the use of drugs that block tissue regeneration and repair, and thus lead to an increase in deviation D. Example: gastrointestinal damage associated with the use of COX inhibitors ^158^.

Even though the pathologic consequences of these different scenarios may be similar, i.e. the loss of a symbiotic microbe or an increase in damage to the host, the ability to distinguish between these pathogenic mechanisms may determine the success of a specific therapy. For example, anti-inflammatory drugs, commonly used to treat IBD, are well adapted to pathogenic scenario 4 characterized by excessive host resistance. However, even though such therapy would increase tissue tolerance to the agent (defined by 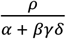 ), it would also decrease resistance to an expanding (scenario 2) or increasingly corrosive (scenario 3) agent, and thereby promote a pathogenic feed-forward loop.

An additional pathogenic scenario derives from a limitation in the host’s resources available for the homeostatic and resistance functions. In that case, inequation 6 is written as:

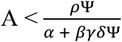

Such limitations would lead to a combination of pathogenic scenarios 4 and 5, and thus to both a loss of control of the microbial agent and decreased tissue regeneration and repair. Interestingly, in the face of such a risk, hot blooded animals can drastically decrease body temperature to conditions of torpor, effectively running the host with a lower energy budget and, at the same time, affecting the agent’s parameters such as its intrinsic growth rate (defined by θ). Fever, the opposite strategy, mobilizes increased levels of energy to fuel the homeostatic and resistance functions, however at the risk of exhausting the host ^159^.

